# Developmentally unique cerebellar processing prioritizes self- over other-generated movements

**DOI:** 10.1101/2023.12.16.571990

**Authors:** Angela M. Richardson, Greta Sokoloff, Mark S. Blumberg

## Abstract

Animals must distinguish the sensory consequences of self-generated movements (reafference) from those of other-generated movements (exafference). Only self-generated movements entail the production of motor copies (i.e., corollary discharges), which are compared with reafference in the cerebellum to compute predictive or internal models of movement. Internal models emerge gradually over the first three postnatal weeks in rats through a process that is not yet fully understood. Previously, we demonstrated in postnatal day (P) P8 and P12 rats that precerebellar nuclei convey corollary discharge and reafference to the cerebellum during active (REM) sleep when pups produce limb twitches. Here, recording from a deep cerebellar nucleus (interpositus, IP) in P12 rats of both sexes, we compared reafferent and exafferent responses to twitches and limb stimulations, respectively. As expected, most IP units showed robust responses to twitches. However, in contrast with other sensory structures throughout the brain, relatively few IP units showed exafferent responses. Upon finding that exafferent responses occurred in pups under urethane anesthesia, we hypothesized that urethane inhibits cerebellar cortical cells, thereby disinhibiting exafferent responses in IP. In support of this hypothesis, ablating cortical tissue dorsal to IP mimicked the effects of urethane on exafference. Finally, the results suggest that twitch-related corollary discharge and reafference are conveyed simultaneously and in parallel to cerebellar cortex and IP. Based on these results, we propose that twitches provide opportunities for the nascent cerebellum to integrate somatotopically organized corollary discharge and reafference, thereby enabling the development of closed-loop circuits and, subsequently, internal models.

**Significance Statement:** For adult animals to produce flexible and adaptive movements, they must distinguish self- from other-generated movements and learn to anticipate how their body moves through space. The computations required for this capacity occur within the cerebellar system. In early infancy, these computations are not yet established and must develop through sensorimotor experience. Here, we found that self-generated movements—particularly those movements occurring during active sleep—are a preferred source of that sensorimotor experience to the infant cerebellum. This preference appears to be unique to the infant cerebellum and helps us understand how that structure establishes its neural circuitry and functions.

## Introduction

The cerebellum’s ability to distinguish reafference from exafference rests on the fact that only reafference, because it arises from self-generated movement, is accompanied by corollary discharge (Blakemore et al., 2000; Crapse and Sommer, 2008; Cullen, 2004). Information from reafference and corollary discharge is used to compute internal models of movement that, in turn, allow animals to discriminate expected from unexpected stimuli (Miall et al., 1993; Wolpert et al., 1998; Bastian, 2006; Person, 2019). Given that the cerebellum exhibits protracted postnatal development (Altman, 1969; Kano et al., 2018) and that internal models must accurately reflect the specific features of individual bodies as they grow and develop, it follows that postnatal sensorimotor experience guides the emergence and recalibration of internal models. But what kinds of sensorimotor experiences are available to infant animals? Noting that active (or REM) sleep is the predominant behavioral state of early infancy (Blumberg et al., 2013), we proposed that the abundant and discrete self-generated limb twitches that characterize that state are better suited to building internal models than wake movements (Mukherjee et al., 2018; Dooley et al., 2021).

In early development, limb movements are produced by brainstem motor structures that project to motor neurons in the spinal cord. In addition, these brainstem motor structures project to precerebellar nuclei, including the inferior olive (IO) and lateral reticular nucleus (LRN; Mukherjee et al., 2018; Alstermark and Ekerot, 2015). As shown in postnatal day (P) 8 rats, neurons in the IO and LRN receive motor-related signals but are not themselves involved in the production of movement (Mukherjee et al., 2018), thus satisfying key criteria of a corollary discharge (Poulet and Hedwig, 2007; Sommer and Wurtz, 2008). Accordingly, at this early age, corollary discharges are conveyed via the IO and LRN to the cerebellar cortex and interpositus nucleus (IP) (Sokoloff et al., 2015a; Sokoloff et al., 2015b; Del Rio-Bermudez et al., 2016). These aspects of IO and LRN function may be specific to the early postnatal period: In adults, in which there currently is no evidence that corollary discharge drives IO activity, there is ample evidence that exafference does (Gibson et al., 2004; Heffley et al., 2018; Ju et al., 2019).

In our earlier studies we did not compare cerebellar activity in response to twitches and manual stimulations; such comparisons are important for assessing the significance of corollary discharge for the developing cerebellum before the onset of a functionally precise internal model. Therefore, here we recorded extracellular activity in the IP of P12 rats, an age when IP exhibits robust responses to twitches (Del Rio-Bermudez, 2016), to assess its ability to distinguish input associated with twitches and stimulations. Because rats around this age reliably exhibit robust responses to stimulation in sensorimotor structures across the neuraxis (Gómez et al., 2023; Gómez et al., 2021; Tiriac and Blumberg, 2016; Del Rio-Bermudez et al., 2015), we expected to see similar responses in IP. Contrary to that expectation, however, exafferent responses in IP were infrequent and weak. Next, we discovered serendipitously that when pups were anesthetized with urethane, robust exafferent responses emerged. Based on this result, we hypothesized that urethane exerted its effect by inhibiting cerebellar cortex, thereby disinhibiting IP. When we tested this hypothesis by ablating the cerebellar cortex, we also found that stimulations evoked strong IP responses and that the responses to twitches were unaffected. Finally, in a separate cohort of P12 rats, we recorded from Purkinje cells in cerebellar cortex and found that their twitch-related activity profiles are nearly identical to those of IP units, suggesting that corollary discharge acts within IP itself to modulate that structure’s response to reafference.

Altogether, these findings suggest a model of cerebellar function at this age in which input arising from self-generated movements during active sleep is prioritized over that from other-generated movements. By providing opportunities for the infant cerebellum to receive converging input from corollary discharge and reafference, such prioritization may be necessary for the development of somatotopically organized closed-loop circuits (Najac and Raman, 2017).

## Materials and Methods

All experiments were conducted in accordance with the National Institutes of Health Guide for the Care and Use of Laboratory Animals (NIH Publication No. 80–23) and were approved by the Institutional Animal Care and Use Committee of the University of Iowa.

### Subjects

Male and female Sprague-Dawley rats (48 total, 25 female) at P12-13 (hereafter P12; body weight: 31.34 ± 3.6 g) were used. Pups were born to dams housed in standard laboratory cages (48 × 20 × 26 cm) with a 12-hr light/dark cycle. Food and water were available ad libitum. The day of birth was considered P0 and litters were culled to eight pups by P3. Pups were randomly assigned to different experimental groups and, to protect against litter effects, pups selected from the same litter were never assigned to the same experimental group (Abbey and Howard, 1973; Lazic and Essioux, 2013).

### Surgery

Surgery was performed using established methods (Blumberg et al., 2015; Glanz et al., 2021). Briefly, on the day of testing a pup with a healthy body weight and a visible milk band was removed from the litter. Under isoflurane anesthesia (3.5–5%, Phoenix Pharmaceuticals, Burlingame, CA), bipolar electrodes (California Fine Wire, Grover Beach, CA) were inserted bilaterally into the nuchal muscle and secured with collodion (Avantor Performance Materials, LLC; Rachor, PA). For those pups receiving subcutaneous infusions of urethane (n = 8; 5 female) or saline (n = 8; 4 female), a small incision (∼1 cm) was made in the skin near the base of the tail and surgical-grade silicon tubing (inner diameter: 0.020 in; outer diameter: 0.037 in; SAI Infusion Technologies, Lake Villa, IL) was inserted subcutaneously and secured with veterinary adhesive (Vetbond, 3M, St. Paul, MN).

For all surgeries, an anti-inflammatory agent (Carprofen, 0.1 mg/kg SC; Putney, Portland, ME) was administered and the torso of the pup was wrapped in soft surgical tape (Micropore, 3M). The scalp was sterilized with iodine and isopropyl alcohol, and a portion of the scalp was removed to reveal the skull; a topical analgesic (bupivacaine, 0.25%; Pfizer, New York, NY) was applied to the skull surface and surrounding skin, and Vetbond was used to secure the skin to the skull. A stainless-steel head-fix (Neurotar, Helsinki, Finland) was attached to the skull using cyanoacrylate adhesive (Loctite, Henkel Corporation, Westlake, OH) that was dried with accelerant (Insta-Set, Bob Smith Industries, Atascadero, CA).

Next, while still under anesthesia, the pup was then secured in a stereotaxic apparatus (Kopf Instruments, Tujunga, CA) and a steel trephine (1.8 mm; Fine Science Tools, Foster City, CA) was used to drill openings in the skull. For interpositus recordings, a hole was drilled over right side of the cerebellum to record from the anterior interpositus nucleus (IP; coordinates from lambda: −1.8 mm AP; +2.0 mm ML). For those pups with cerebellar cortical ablations (n = 8; 4 female), the right portion of the interparietal bone was trephined to expose and disconnect the right cerebellar cortex from the deep cerebellar nuclei. This procedure entailed either ablation of cortical tissue using a surgical hook and forceps or disconnection (without tissue removal) using a 27G needle (BD Precision Glide; Franklin Lakes, NJ), bent at a 90-degree angle. For sham pups (n = 8; 3 female), only the interparietal bone was removed. Finally, for recordings in cerebellar cortex (n = 16 pups; 9 female; 6 pups were excluded due to loss of units or lack of forelimb responsivity to twitches), the procedure was similar to that described above, but the electrode was placed in the cortex (right crus I/II) immediately dorsal to IP (coordinates from lambda: 2.3 +/- 0.3 mm AP; 2.2 +/- 0.1 mm ML).

Throughout surgery, the pup’s respiration was monitored and the concentration of isoflurane was adjusted accordingly. All surgical procedures lasted 16-24 min. After surgery was complete, the pup was transported to the recording rig, where it was head-fixed in a warm environment for at least 1 h. Recording began only after regular sleep– wake cycles were observed and intracranial temperature was at least 36°C.

### Data Acquisition

To record neural activity, a 16-site linear silicon electrode (100 µm between sites; NeuroNexus, Ann Arbor, MI) was lowered into IP (−3.3 +/- 0.3 mm DV) or cerebellar cortex (−2.5 +/- 0.5 mm). Before insertion, the electrode was coated with fluorescent DiI (Life Technologies, Carlsbad, CA) to enable histological confirmation of electrode location. Neurophysiological recordings were accomplished using a data acquisition system (Tucker-Davis Technologies, Alachua, FL). EMG and neural activity were acquired at 1 kHz and 25 kHz, respectively. Video was acquired at 100 frames/s using a BlackFly-S camera (FLIR Integrated Systems, Wilsonville, OR) and recorded using SpinView software (FLIR). Video and neural data were synchronized using a custom MATLAB script as described previously (Dooley et al., 2021).

#### IP activity preceding and following urethane injection

Neurophysiological, EMG, and video data were recorded for 60 min as the pup cycled freely between sleep and wake. After 60 min, ∼50 manual stimulations of the right forelimb, ipsilateral to the recording site, were delivered using a cotton-tipped wooden dowel so as to rapidly displace the limb (Gómez et al., 2021). Stimulations were delivered at random intervals, 5-15 s apart and at least 2 s after the cessation of a limb movement, during active sleep or wake; stimulations that evoked a movement were excluded from analysis. To estimate the duration of the stimulations (i.e., the time from the onset of the application of the dowel to its removal), we sampled 10 stimulations from 10 pups (i.e., 100 total) and calculated a mean duration of 339 ± 131 ms. After the stimulations were delivered, urethane (1.0 mg/g b.w.; Sigma-Aldrich, St. Louis, MO) or an equivalent volume of sterile saline (1.0 ml/g b.w.; Hospira Inc, Lake Forest, IL) was administered via the subcutaneous cannula. After 15 min, another set of ∼50 stimulations of the right forelimb were delivered as described above.

#### IP activity after ablation of cerebellar cortex

Data were collected and stimulations were delivered as described above. Following stimulations, recordings continued for an additional 45 min to collect data as pups cycled freely between sleep and wake.

#### Cerebellar cortical activity

Neurophysiological, EMG, and video data were recorded for 60 min as the pup cycled freely between sleep and wake.

### Histology

At the end of the recording period, the pup was euthanized with 0.05 ml/kg of ketamine-xylazine (90:10; 0.002 ml/g b.w. IP Akorn, Inc.; Lake Forest, IL) and perfused transcardially with phosphate-buffered saline (PBS) followed by 4% paraformaldehyde (PFA). The brain was extracted from the skull and placed in PFA for at least 24 h, after which it was transferred to phosphate-buffered sucrose for at least 48 h before sectioning. The cerebellum was sectioned coronally at 80 µm. Electrode locations were determined using fluorescent microscopy before Nissl staining.

### Data Analysis

#### Spike sorting

Neurophysiological, EMG, and behavioral data were imported to MATLAB and analyzed as described previously (Gómez et al., 2023). Briefly, we used custom scripts to filter raw neurophysiological data (band-pass: 500-5,000 Hz) and extracted units using Kilosort (Pachitariu et al., 2016); subsequent unit templates were visualized in Phy2 (Rossant & Harris, 2022). Units in cerebellar cortex were analyzed using template matching in Spike2 (cerebellar cortex; Sokoloff et al., 2015a); we restricted this analysis to Purkinje cells (38 of a total of 66 twitch-responsive units) identified by their waveform and spike amplitude (Crepel, 1971; Sokoloff et al., 2015b). In IP, only data from single units were analyzed; in cerebellar cortex, both single units and multi-units were analyzed.

#### Video analysis of movement and behavioral-state determination

Synchronized electrophysiological and behavioral data were imported into Spike2 (Cambridge Electronic Design, Cambridge, UK). Based on EMG activity and behavior (from video), Spike2 was used to define behavioral states, as described previously (Del Rio-Bermudez et al., 2016). Periods of high muscle tone and atonia were identified when nuchal EMG was above or below, respectively, an average threshold of 3x baseline (Gómez et al., 2023). Movements of the limb were detected using a custom MATLAB script that identified movement based on frame-by-frame changes in pixel intensity within regions-of-interest (ROIs; Dooley et al., 2020). Using an ROI encompassing the right forelimb, the number of pixels with >5% changes in pixel intensity were summed frame by frame, resulting in a trace of real-time movement. Using a threshold for displacement of the right forelimb, events were triggered at movement onset, and subsequently confirmed visually by the experimenter using video. Finally, the onset of a limb stimulation was scored manually based on the frame immediately preceding visual displacement of the limb. Manual stimulations that evoked sudden increases in nuchal EMG and large movements of multiple limbs were not included in the analysis.

Active wake was defined as a period of high muscle tone when the pup was also moving its limbs in a coordinated fashion. Active sleep was defined as a period of muscle atonia punctuated by sharp spikes in the EMG record and rapid changes in pixel intensity, indicative of myoclonic twitching. The remaining periods were defined as behavioral quiescence and were not analyzed further.

#### Determination of responsive and nonresponsive units

We characterized units in IP and cerebellar cortex either responsive or nonresponsive to twitch-related reafference or stimulation-related exafference. First, baseline firing rate was calculated for each unit based on its activity 250-500 ms before movement onset. Second, a unit was considered responsive to a twitch or stimulus if peak mean firing rate for the unit within 250 ms of movement onset exceeded 3.5 standard deviations above baseline firing rate.

#### Width at half-height

To determine width at half-height, neural data were normalized to baseline activity (baseline = 0; peak = 1). Data were then smoothed using a 5-bin kernel before being interpolated from 10-ms bins to 1-ms bins (using the MATLAB interp1() function). Width at half-height was measured as the duration (in ms) at the point halfway between baseline (0) and peak (1).

### Statistical Analysis

To compare the proportion of responsive units before and after the injection of urethane or saline in the IP, a within-subjects repeated-measures analysis of variance (ANOVA) was conducted. A Shapiro-Wilk test for normality and a Levene’s test for homogeneity were performed on all comparisons before analysis; in the event a comparison failed either of these tests, Welch’s t test was used instead. To compare the proportion of responsive units following cerebellar cortical ablation, a between-subjects one-way ANOVA was performed. Post-hoc analyses were performed when appropriate. All statistical analysis of the proportion of responsive and nonresponsive neurons was performed using an arc-sin transformation. A chi-squared test was used to determine if the proportion of neurons that responded to twitches in unanesthetized animals were the same as those that responded to exafference following urethane administration. Between-subject t tests were performed to compare average width-at-half-height, and peak response time, and amplitude between stimulus responsive units.

All statistical tests were performed using SPSS (IBM; Chicago, IL). Unless otherwise noted, means are presented with standard errors and alpha was set at 0.05. Effect size was calculated using an adjusted partial eta-squared for ANOVAs (adj η_p_^2^) and adjusted eta-squared for t tests (adj η^2^), adjusted against positive bias (Mordkoff, 2019). When appropriate, a Bonferroni correction was used to correct alpha for multiple comparisons.

## Results

To distinguish differences in IP responses to reafference and exafference, we recorded extracellular unit activity in head-fixed P12 rats (**Figure 1A**). For each pup, we recorded 60 min of spontaneous sleep-wake cycling, followed by ∼50 manual stimulations of the forelimb ipsilateral to the recording site (across pups, 36-48 stimulations met criteria; **Figure 1B**). In prior work at the same ages using similar methods, we did not observe effects of behavioral state on neural responsivity to stimulation (Gómez et al., 2021; Tiriac et al., 2016; therefore, stimulations were performed here during both sleep and wake when pups were not moving. After completion of the stimulation protocol, urethane or saline was administered and, after 20 min, the forelimb was stimulated again (40-50 stimulations/pup met criteria). Electrode location in IP was confirmed histologically (**Figure 1C**). In pups injected with urethane, we recorded 78 units (n = 8 pups; 8-17 units/pup); in pups injected with saline, we recorded 86 units (n = 8 pups; 7-22 units/pup).

**Figure 1.**
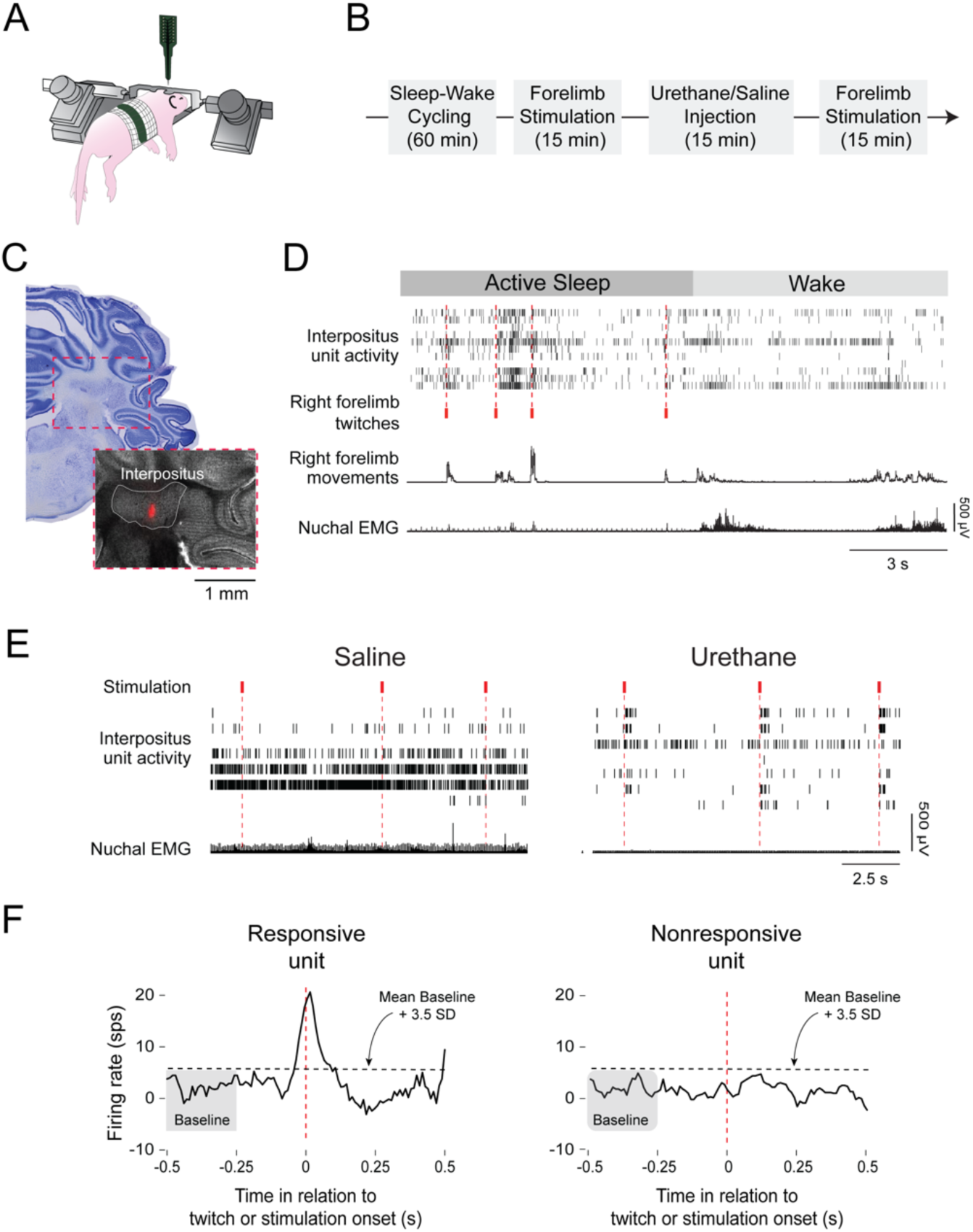
Experimental design and representative data. **A**, Illustration showing the head-fixed pup in the recording apparatus. **B**, Timeline of the experimental procedure. ***C***, Representative histology showing the red-florescent electrode track in interpositus (inset). ***D***, Representative data from recording in a P12 rat. From top: Behavioral states marked as active sleep (dark gray) or wake (light gray), black ticks showing interpositus spiking activity for individual units in each row, red ticks showing identified twitches of the right forelimb, movements of the right forelimb derived from video, and nuchal EMG activity. ***E***, Representative data from pups after saline (left) or urethane (right) injection. From top: Red ticks denoting the onset of a forelimb stimulations, black ticks showing interpositus spiking activity for individual units in each row, and nuchal EMG activity. ***F***, Representative units illustrating method for identifying units that were responsive (left) and nonresponsive (right) to twitches or stimulations. See Methods for additional detail.

A representative recording of IP unit activity, right forelimb movements, scored twitches, and nuchal EMG during active sleep and wake is shown in **Figure 1D**. The recording illustrates four forelimb twitches and IP activity surrounding those events. Two additional representative recordings during the stimulation period are shown for pups after administration of saline or urethane (**Figure 1E**). Three forelimb stimulations are shown for each condition; for the urethanized pup, the increase in IP activity after each stimulation is apparent. Twitches and stimulations were used as triggers for perievent histograms of unit firing rate that, in turn, were used to assess whether a unit was responsive or nonresponsive (**Figure 1F**; see Methods).

### IP units are more likely to respond to twitches than to stimulations

As found previously in P12 rats (Del Rio-Bermudez et al., 2016), most IP neurons exhibited pronounced increases in activity in the period surrounding a twitch (this was true in both the pre-saline and pre-urethane pups; **Figures 2A-B**). In contrast, IP neurons were less likely to respond to stimulation. The proportion of responsive units for twitches was approximately 3x that for stimulations (70-76% vs. 25-22%; *p*s < .001, binomial test).

**Figure 2.**
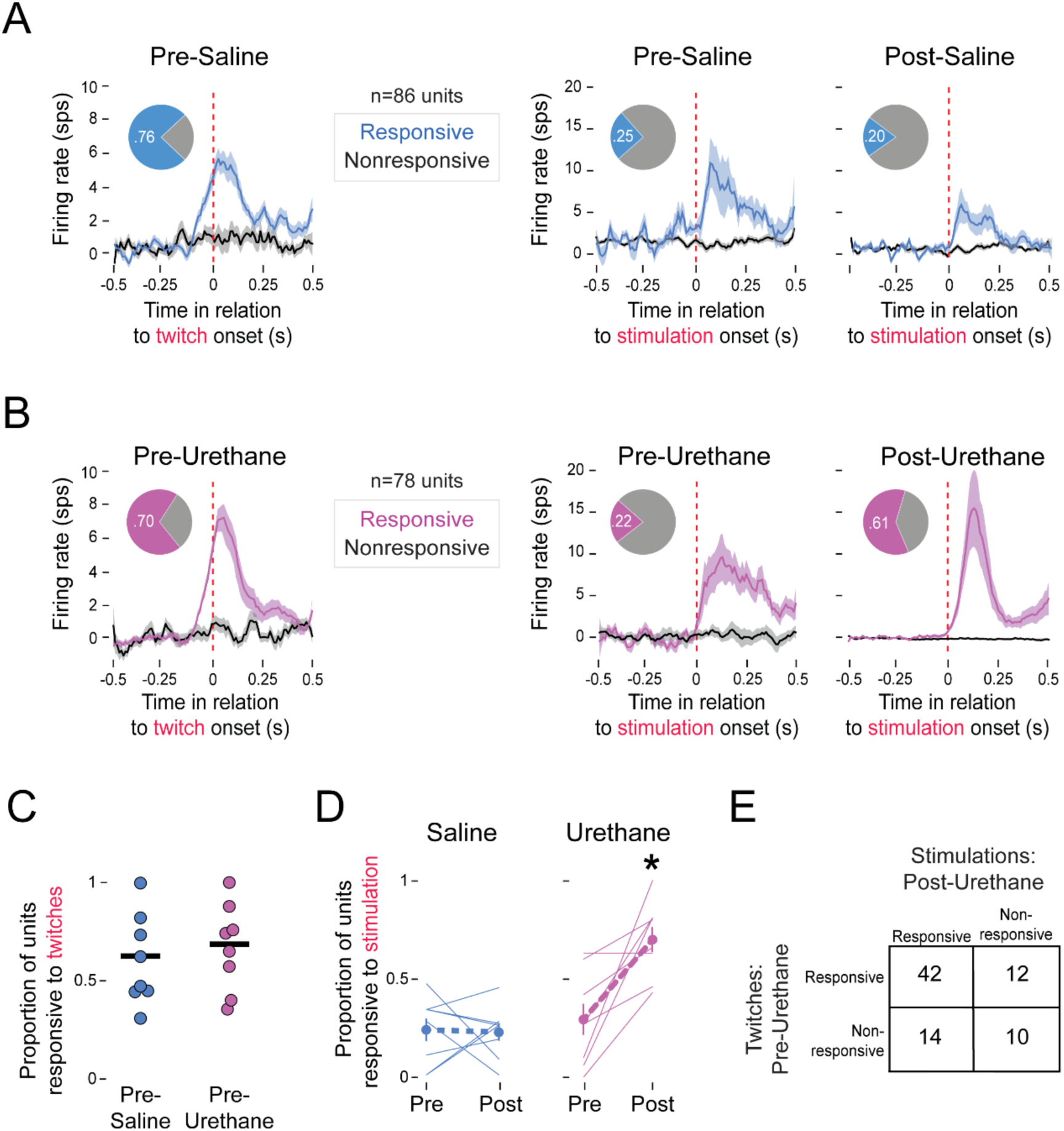
Responsiveness of interpositus units to twitches and stimulations before and after urethane or saline injection. ***A***, Perievent time histogram of unit firing rate for responsive (blue) and nonresponsive (black) units in the interpositus in relation to the onset of forelimb movements associated with twitches or stimulations (red dotted lines) before and after saline injection. Firing rates are normalized to baseline. Pie charts represent proportion of responsive and nonresponsive units. Note the different y-axis ranges. ***B***, Same as in ***A*** but for pups before and after urethane injections (responsive units: magenta). ***C***, Mean proportion of twitch-responsive units for each pup before injection of saline or urethane. n = 8 pup/group. ***D***, Mean proportion of stimulation-responsive units before and after saline (left) or urethane (right) injection. Asterisk denotes significant difference in relation to the post-saline pups. ***E***, Latin square showing the association between IP units that were or were not responsive to twitches (pre-urethane) and stimulations (post-urethane). Units that were responsive to twitches were significantly more likely to also be responsive to stimulations, *X^2^* = 35.03, *p* < 0.001.

### IP units respond to exafference after urethane anesthesia

After recording IP responses to reafference and exafference, pups received either saline or urethane before the stimulation protocol was repeated (note that urethane suppresses twitches). Pooling units across pups, there was no difference in the proportion of stimulation-responsive units before and after administration of saline (**Figure 2A**). In contrast, urethane administration significantly increased the proportion of stimulation-responsive units (20% vs. 61%; *p* < .0001, binomial test; **Figure 2B**). Similarly, there was no significant difference in the proportion of twitch-responsive units per pup (*t*_(14)_ = 0.628; **Figure 2C**); for stimulation-responsive units, there was a significant main effect of group (urethane vs. saline; *F*_(1,14)_ = 15.93, *p* = 0.001, adj *η_p_^2^* = 0.499) but not time (pre vs. post; *F*_(1,14)_ = 4.41), and a significant group x time interaction (*F*_(1,14)_ = 6.53, *p* = 0.023, adj *η_p_^2^* = 0.269; **Figure 2D**). A post hoc test confirmed that the proportion of responsive IP units was significantly higher in the Post-Urethane group (t_(1,14)_ = 43.36, *p* < 0.001, adj *η^2^* = 0.738).

Because pups’ limbs dangle freely in our apparatus, twitches trigger predominantly proprioceptive input; in contrast, because stimulations involve externally produced movements of the limb, they trigger both proprioceptive and tactile input (Gómez et al., 2021). Given this overlapping engagement of proprioception, we expected to see an association between those IP units that were responsive to twitches in the Pre-Urethane group and those units that were responsive to stimulations in the Post-Urethane group. Indeed, there was a significant association, with units exhibiting dual responsivity to twitches and stimulations being more prevalent than all other possibilities combined (*X^2^* = 35.03, *p* < 0.001; **Figure 2E**).

### Reafferent and exafferent IP responses exhibit different response profiles

Why were twitches more effective than stimulations in triggering IP responses? The most likely explanation relates to the fact that twitches, as self-generated movements, are accompanied by corollary discharges (Mukherjee et al., 2018). Accordingly, we determined whether corollary discharge is an observable component of the twitch-related profile of IP units. To do this, we compared twitch-related unit responses (in the Pre-Urethane group) and stimulation-related unit responses (in the Post-Urethane group; **Figure 3A-B**); the latter presumably comprise only a sensory response. In this comparison of response profiles, we focused on the width at half-height (a measure of the duration of the response profile) and the timing of the response peak (which provides insight into the whether the peak reflects sensory feedback, corollary discharge, or a combination of the two).

**Figure 3.**
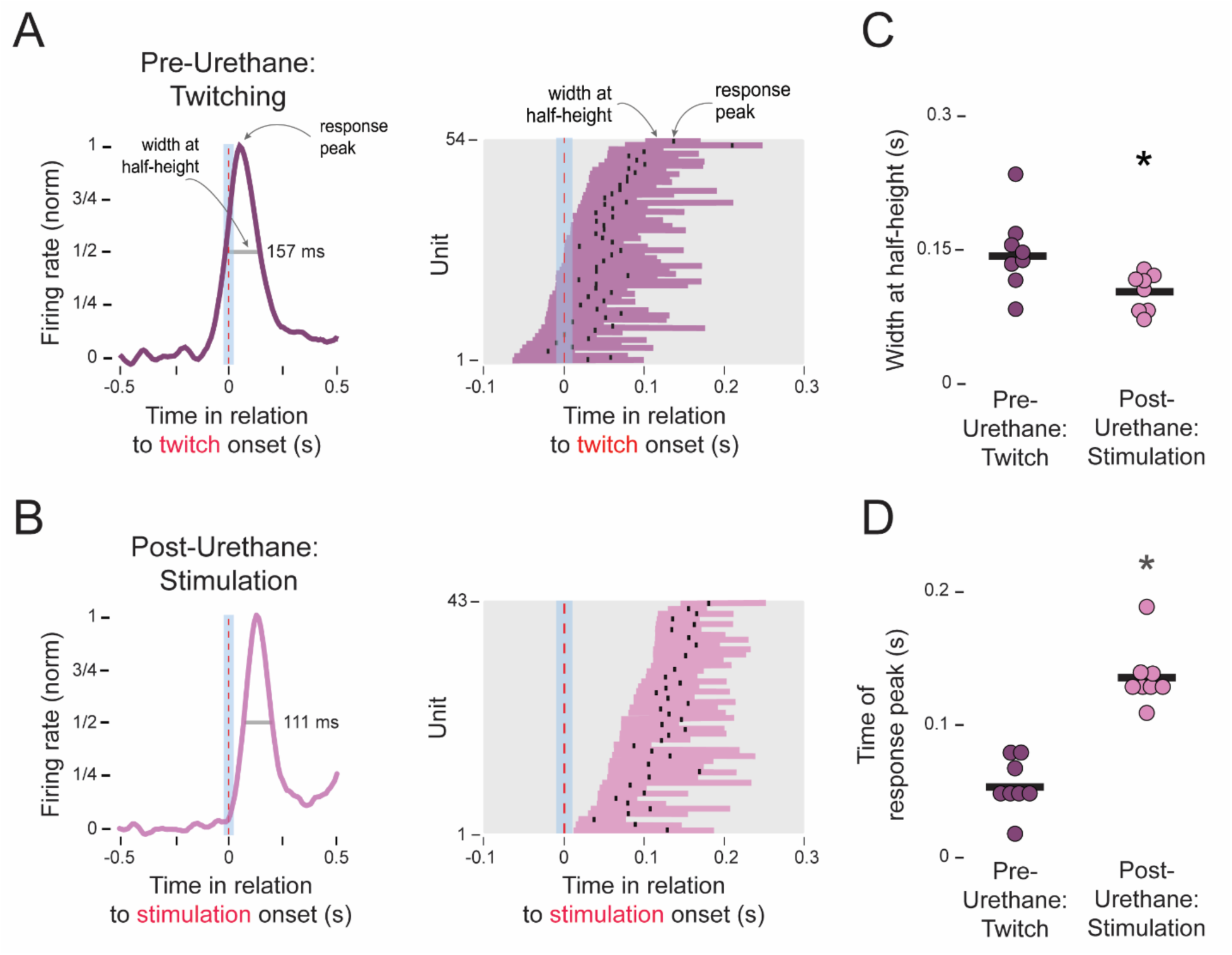
Interpositus response profiles for twitches and stimulations. ***A***, Left: Representative perievent time histogram for a twitch-responsive unit (pre-urethane). The histogram is normalized to baseline and peak amplitude. Right: Representations of the range of width at half-height for all individual twitch-responsive units. Response peaks are also shown as black ticks. ***B***, Same as in ***A*** but for stimulation-responsive units (post-urethane). ***C***, Mean width-at-half-height for twitch- and stimulation-responsive units. n = 8 pups/group. Asterisk denotes significant difference between groups. ***D***, Same as in ***C*** but for mean time of response peak.

Compared with the IP unit response profiles for stimulations, the twitch-related profiles exhibited several distinct features: They were typically initiated earlier, had significantly longer-duration widths at half-height (*t*_(14)_ = 2.49, *p* = 0.031, adj *η^2^* = 0.303; **Figure 3C**), and had a significantly earlier peak response (*t*_(14)_ = 7.52, *p* < 0.001, adj *η^2^* = 0.900; **Figure 3D**). Together, these findings suggest that the combined inputs of corollary discharge and reafference are detectable in the response profiles of IP units at this age.

### Ablation of cerebellar cortex enables exafference in IP

We hypothesized that urethane’s enabling of IP responses to exafference resulted from inhibition of Purkinje cells, thereby decreasing inhibitory outflow from the cortex. IP’s responsivity to twitch-related reafference might also result from a release of cortical inhibition, but the mechanism of that release might rely on the action of corollary discharge—acting within cerebellar cortex or IP itself.

To assess the functional contributions of cerebellar cortex to IP responses to twitches and stimulations, we ablated cerebellar cortex in P12 rats and subsequently recorded ipsilateral IP activity (**Figures 4A-B**). We recorded from 65 units in pups with cortical ablations (n = 8 pups; 6-11 units/pup) and 66 units in pups with sham ablations (n = 8 pups; 6-14 units/pup; **Figure 4C-D**). Whereas cortical ablation had no significant effect on the proportion of twitch-responsive units (*t*_(14)_ = 1.29; **Figure 4E**), it significantly increased the proportion of stimulation-responsive units (from 25% to 80% of units; *t*_(14)_ = 25.74, *p* < 0.001, adj *η^2^* = 0.623; **Figure 4F**). This increase in exafferent responsivity after cortical ablation supports the notion that tonic cortical inhibition of IP normally suppresses exafference at this age.

**Figure 4.**
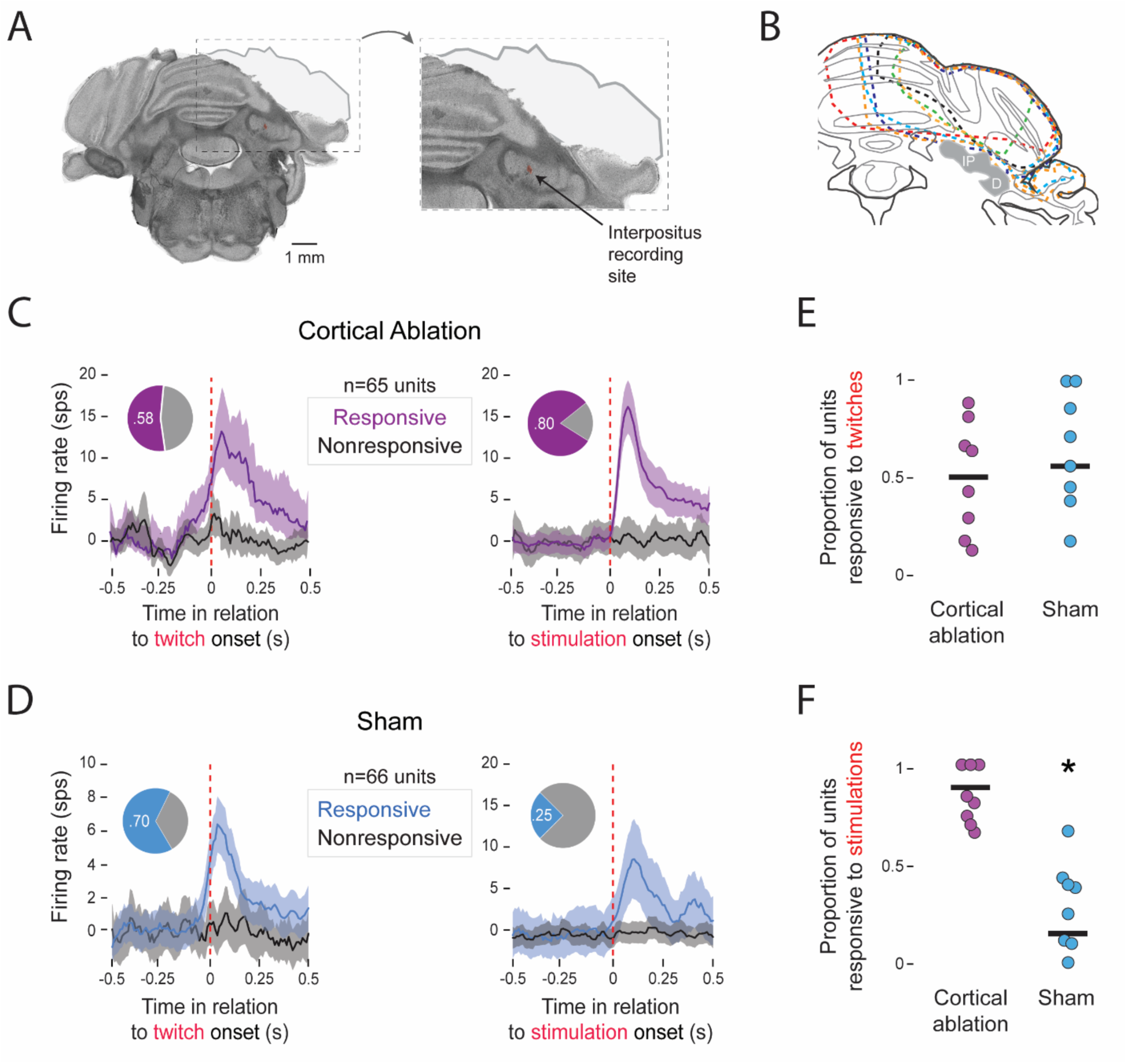
Effect of cortical ablation on responsiveness of interpositus units to twitches and stimulations. ***A***, Representative coronal section showing unilateral ablation of cerebellar cortex. Inset shows location of electrode in interpositus (red-fluorescent dot). ***B***, Extent of cerebellar cortical ablations in individual pups denoted by dotted lines of different colors. IP, interpositus nucleus; D, dentate nucleus. ***C***, Perievent time histogram of unit firing rate for responsive (violet) and nonresponsive (black) units in the interpositus in relation to the onset of forelimb movements associated with twitches or stimulations (red dotted lines) in pups with cortical ablations. Firing rates are normalized to baseline. Pie charts represent proportion of responsive and nonresponsive units. ***D***, Same as ***C*** but for pups with sham ablations. Note the different y-axis ranges. ***E***, Mean proportion of twitch-responsive units for pups with cortical ablations and sham ablations. n = 8 pups/group. ***F***, Same as ***E*** but for stimulation-responsive units. Asterisk denotes significant difference between groups.

### Ablation of cerebellar cortex does not affect reafference

The fact that cortical ablation did not affect the ability of IP units to exhibit twitch-related responses indicates that the cortex is not necessary for those responses. However, it is possible that the cortex asserts more subtle effects on IP responding. We assessed this possibility by comparing the response profiles of IP units during twitching in pups with cortical ablations and sham ablations. Cortical ablation did not significantly affect width at half-height (*t*_(14)_ = 0.10; **Figure 5B**) or time of response peak (Welch’s *t*_(9.28)_ = 5.86; **Figure 5C**). We did observe a significant increase in peak amplitude after cortical ablation (Welch’s *t*_(1,37.33)_ = 3.03, *p* = 0.004, adj *η^2^* = 0.050; **Figure 5D**), but this difference was not significant across pups (Welch’s *t*_(1,8.55)_ = 0.81; **Figure 5E**). Cortical ablation did produce significant increases in the variance of the time of peak response (Levene’s *F*_(1,14)_ = 9.49, *p* = 0.008, adj *η_p_^2^* = 0.361) or peak amplitude (Levene’s *F*_(1,14)_ = 6.17, *p* = 0.026, adj *η_p_^2^* = 0.256). These last results suggest that the cerebellar cortex exerts a tonic modulatory influence on IP activity, even if it does not mediate the effect of corollary discharge on IP responsivity.

**Figure 5.**
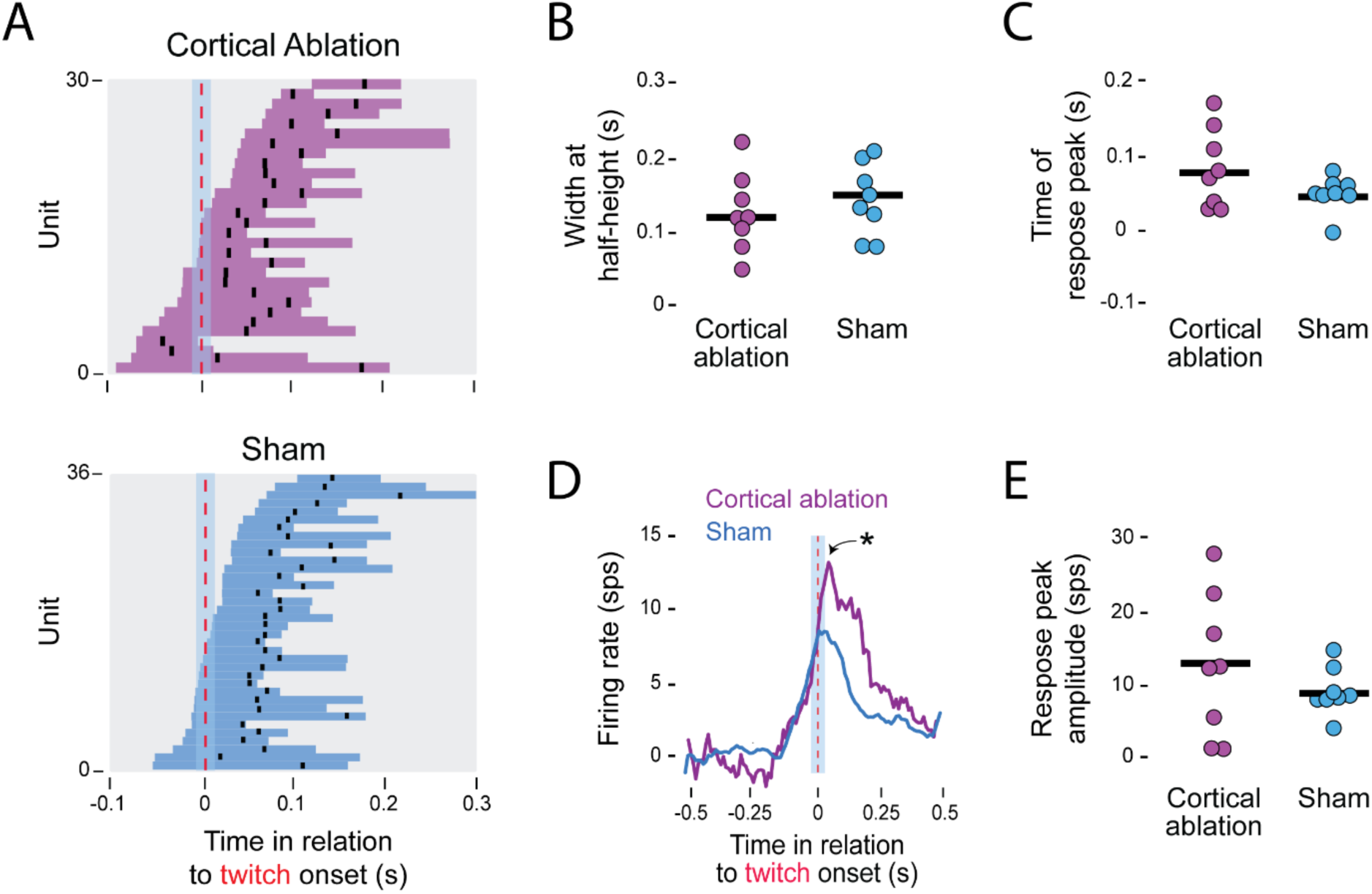
Effect of cortical ablation on interpositus response profiles for twitches. ***A***, Representations of the range of width at half-height for all individual twitch-responsive units for pups with cortical ablations (top) and sham ablations (bottom). Response peaks are also shown as black ticks. ***B***, Mean width-at-half-height for twitch-responsive units in pups with cortical ablations and sham ablations. n = 8 pups/group. ***C***, Same as in ***B*** but for mean time of response peak. ***D***, Perievent time histogram of mean unit firing rate (normalized to baseline) for cortical ablation (n = 30) and sham (n = 36) units pooled across pups. Asterisk indicates significant group difference in peak amplitude. ***E***, Same as in ***B*** but for mean response-peak amplitude.

### Parallel reafferent input to IP and cerebellar cortex

Having demonstrated that IP does not depend on cerebellar cortex for its reafferent response, the question arises as to whether cerebellar cortex receives twitch-related input before, after, or at the same time as IP. We addressed this question by recording activity in cerebellar cortex in a separate cohort of P12 rats. Across 10 pups, a total of 38 twitch-responsive Purkinje cells were identified (1-8 units/pup; **Figure 6A**). We compared the response profiles of these twitch-responsive units to those of the 94 twitch-responsive units in IP (combined across the Pre-Saline and Pre-Urethane groups; n = 94 units from 16 pups; 7-22 units/pup; **Figure 6B**). Whereas width at half-height was significantly reduced in cerebellar cortex (Welch’s *t*_(18.53_) = −5.81, *p* < 0.001, adj *η^2^*= 0.535; **Figure 6C**), there was no significant difference in peak response time (*t*_(24)_ = 0.71; **Figure 6D**). Overlaying the response profiles of twitch-responsive cortical and IP units highlights the near-identical timing of their activity (**Figure 6E**), suggesting that the two structures receive parallel twitch-related input from precerebellar structures at this age.

**Figure 6.**
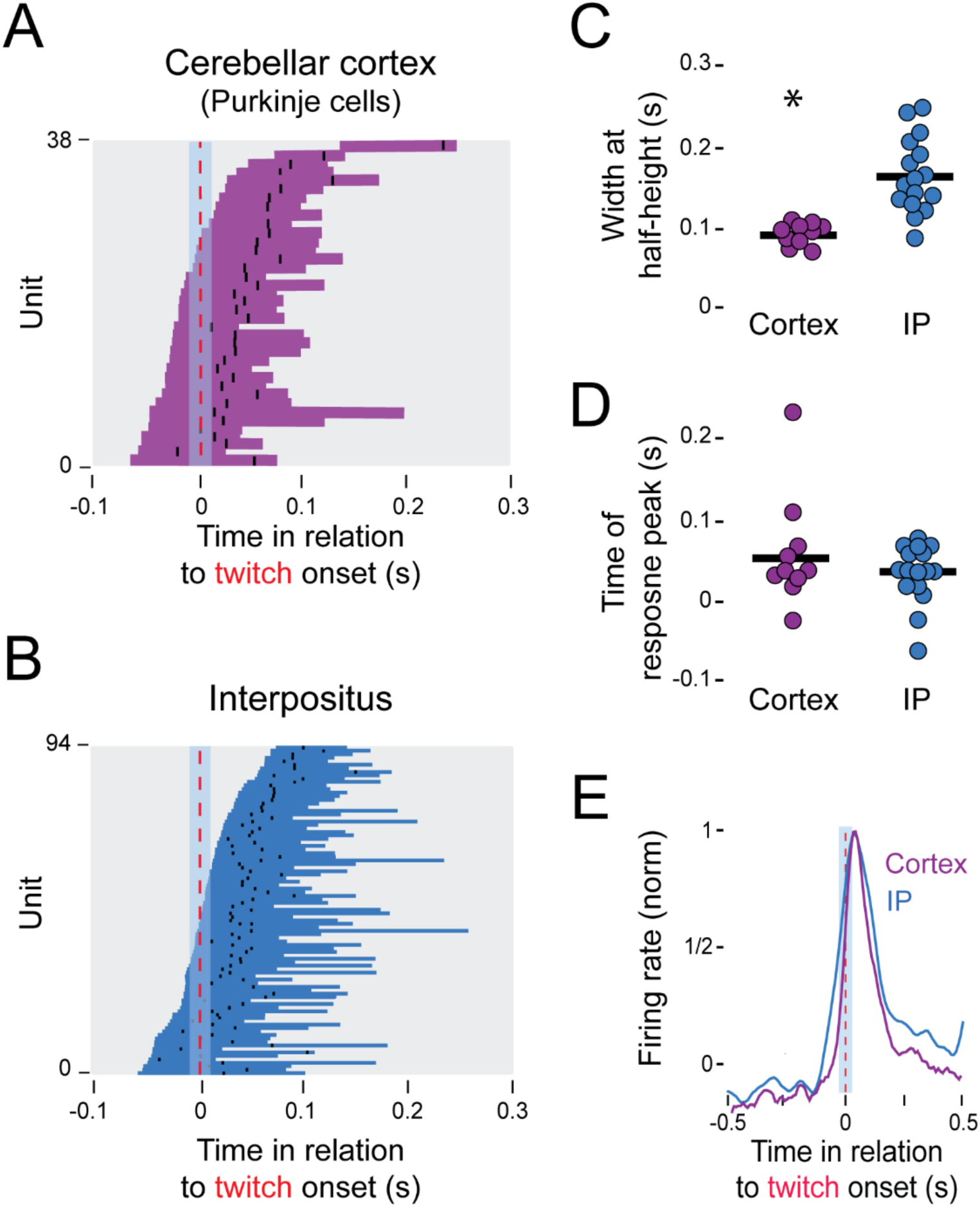
Response profiles for twitch-responsive units in cerebellar cortex and interpositus. ***A***, Representations of the range of width at half-height for all individual twitch-responsive Purkinje cells in cerebellar cortex. Response peaks are also shown as black ticks. ***B***, Same as in ***A*** but for IP (combined across the Pre-Saline and Pre-Urethane groups). ***C***, Mean width-at-half-height for twitch-responsive Purkinje cells in cerebellar cortex (n = 10 pups) and units in interpositus (n = 16 pups). ***D***, Same as in ***C*** but for mean time of response peak. ***E***, Mean firing rate (normalized to baseline and peak amplitude) for twitch-responsive Purkinje cells in cerebellar cortex and units in interpositus.

## Discussion

The infant IP appears to be unique among the sensorimotor structures examined thus far in that it does not readily respond to exafferent input (Gómez et al., 2023; Gómez et al., 2021; Tiriac and Blumberg, 2016; Del Rio-Bermudez et al., 2015). Here we sought to understand the mechanisms responsible for this lack of responsiveness. Serendipitously, we discovered that urethane anesthesia permits the expression of exafference in IP, suggesting that urethane reduces the inhibitory influence from cerebellar cortex. Upon discovering that cortical ablation mimics the effect of urethane, we concluded that the cortex tonically inhibits IP to prevent the expression of exafference (**Figure 7A**). But such a conclusion does not resolve the question of how twitches, unlike stimulations, exert such a powerful activational effect on IP.

**Figure 7.**
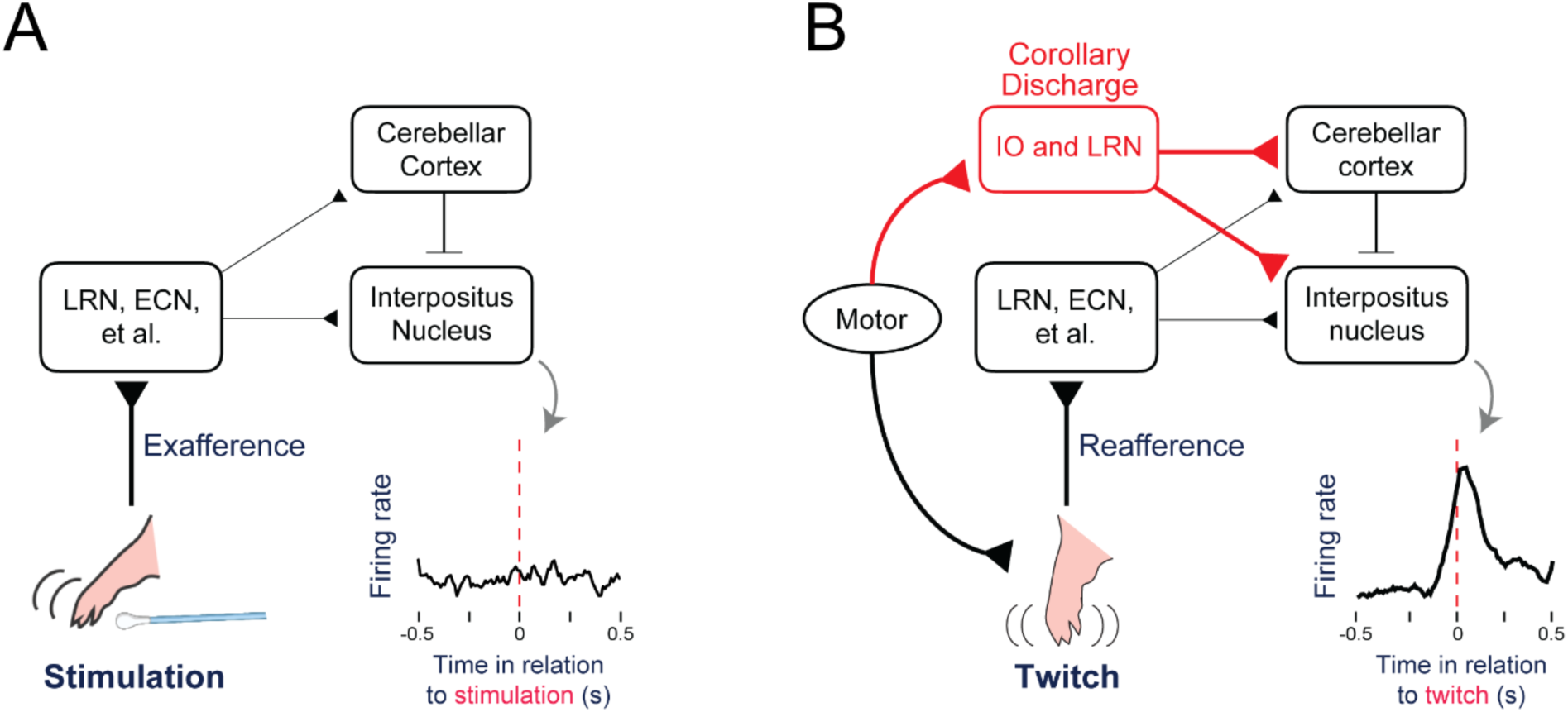
Summary diagrams to explain differential responses of interpositus units to twitches and stimulations in P12 rats. ***A***, When the forelimb is stimulated, exafference is processed in such precerebellar nuclei as the lateral reticular nucleus (LRN) and external cuneate nucleus (ECN) before being conveyed via mossy fibers to interpositus and cerebellar cortex. As shown here, exafference is not expressed in interpositus at this age. ***B***, Twitches are produced by motor commands arising from brainstem premotor nuclei. At the same time, a corollary discharge (red) is conveyed to the inferior olive (IO) or lateral reticular nucleus (LRN). Whereas signals from the IO are sent to the cerebellum via climbing fibers, signals from the LRN are sent via mossy fibers (it is not yet clear which of these two pathways is necessary or sufficient), resulting in a robust IP response that reflects input from both corollary discharge and reafference.

### How are twitches prioritized?

To explain the ability of twitches to preferentially activate the infant IP, we turn to the feature that distinguishes them from stimulations: the production of corollary discharge. In our previous work, we focused on two precerebellar nuclei—the IO and LRN—in part because both receive direct input from motor structures in the mesodiencephalic junction (De Zeeuw et al., 1998; Alstermark and Ekerot, 2013), a region that contains motor nuclei involved in the production of twitching (Del Rio-Bermudez et al., 2015). After corollary discharge is received by the infant IO, it is conveyed to the cerebellum via climbing fibers (Mukherjee et al., 2018; **Figure 7B**; LRN also conveys corollary discharge, but via mossy fibers). In contrast, reafference from twitches and exafference from stimulations are conveyed to the cerebellum via mossy fibers arising from the LRN and other precerebellar nuclei (Alstermark and Ekerot, 2013; Puro and Woodward, 1977). Critically, at P12, climbing fibers provide more powerful cerebellar input than mossy fibers (Najac & Raman, 2017). Accordingly, we propose that the early component of the twitch-related activation of IP—that is, the component associated with corollary discharge—reflects input from climbing fibers that overcomes the relatively weak tonic inhibition from cortical Purkinje cells at this age (Najac & Raman, 2017; Watanabe & Kano, 2011).

But how do we explain the reafferent component of twitch-related IP activity? We had initially hypothesized that corollary discharge from IO activates complex spikes in Purkinje cells, thereby disinhibiting reafference in IP (Davie et al., 2008; Sato et al., 1993; Armstrong and Rawson, 1979). However, this hypothesis has a problem: If corollary discharge causes cerebellar cortex to disinhibit IP activity to allow for the expression of reafference, then Purkinje-cell activity should have decreased as IP activity increased in the period immediately after a twitch. Instead, cortical Purkinje cells and IP units exhibited nearly identical twitch-related increases in activity (**Figure 6E**), a finding that is inconsistent with the former structure modulating the latter’s reafferent responses.

Accordingly, we propose that the expression of twitch-related reafference in IP relies on corollary discharge acting within IP itself to decrease the inhibitory influence of the cerebellar cortex. Such an effect could be mediated at the level of IP interneurons (Chen et al., 2005, Kebschull et al., 2020; Brinke et al., 2017), but whether such a circuit is functional at this age is unclear at this time.

### Why are twitches prioritized?

We are not the first to show that IP receives weak exafference in early development: In P20 but not P12 rats, IP supports classical eye-blink conditioning in which a tone is used as the conditioned stimulus (Campolattaro and Freeman, 2008; Freeman and Muckler, 2003; Stanton et al., 1998). However, conditioning at P12 is possible with direct stimulation of the pons, overcoming the weak exafference from the tone in the developing auditory system (Campolattaro and Freeman, 2008). The investigators concluded that cerebellar neurons in infant animals are able to learn even if they don’t have anything to learn about. However, the abundance of twitch-related corollary discharge and reafference processed by the cerebellum at P12 does suggest that pups are using these signals to learn the relations between motor commands and their sensory consequences.

The infant projections from IO to IP appear to play a critical role in the development of cerebellar circuitry (Lang et al., 2017; Najac and Raman, 2017). These projections, which initially arise from exuberant climbing-fiber collaterals, have a strong excitatory influence on IP that weakens with age (Najac and Raman, 2017; Lu et al., 2016). Consequently, climbing fibers drive much of IP’s activity at a time when the cerebellar system has limited capacity and is still developing its basic functional organization.

Therefore, based on the present results, we propose that the corollary discharge signals conveyed by climbing fibers in early development are sufficiently strong to overcome cortical inhibition and activate IP in ways that exafference alone cannot. Moreover, because the present results suggest that twitch-related corollary discharge and reafference arrives in parallel to the cerebellar cortex and IP, a competitive context is established that involves coincident activity of somatotopically related neurons (**Figure 8**). The parallel activation we observe in the cerebellum at P12 coincides with a known critical period over the second postnatal week during which climbing fiber synapses are substantially pruned (Kano et al., 2018; Hashimoto & Kano, 2013). Accordingly, we hypothesize that twitch-related cerebellar activity shapes the selective pruning of climbing fiber afferents and, more broadly, contributes to the development of functional closed-loop circuits within cerebellar microzones (Sugihara, 2006; Ruigrok and Voogd, 2000).

**Figure 8.**
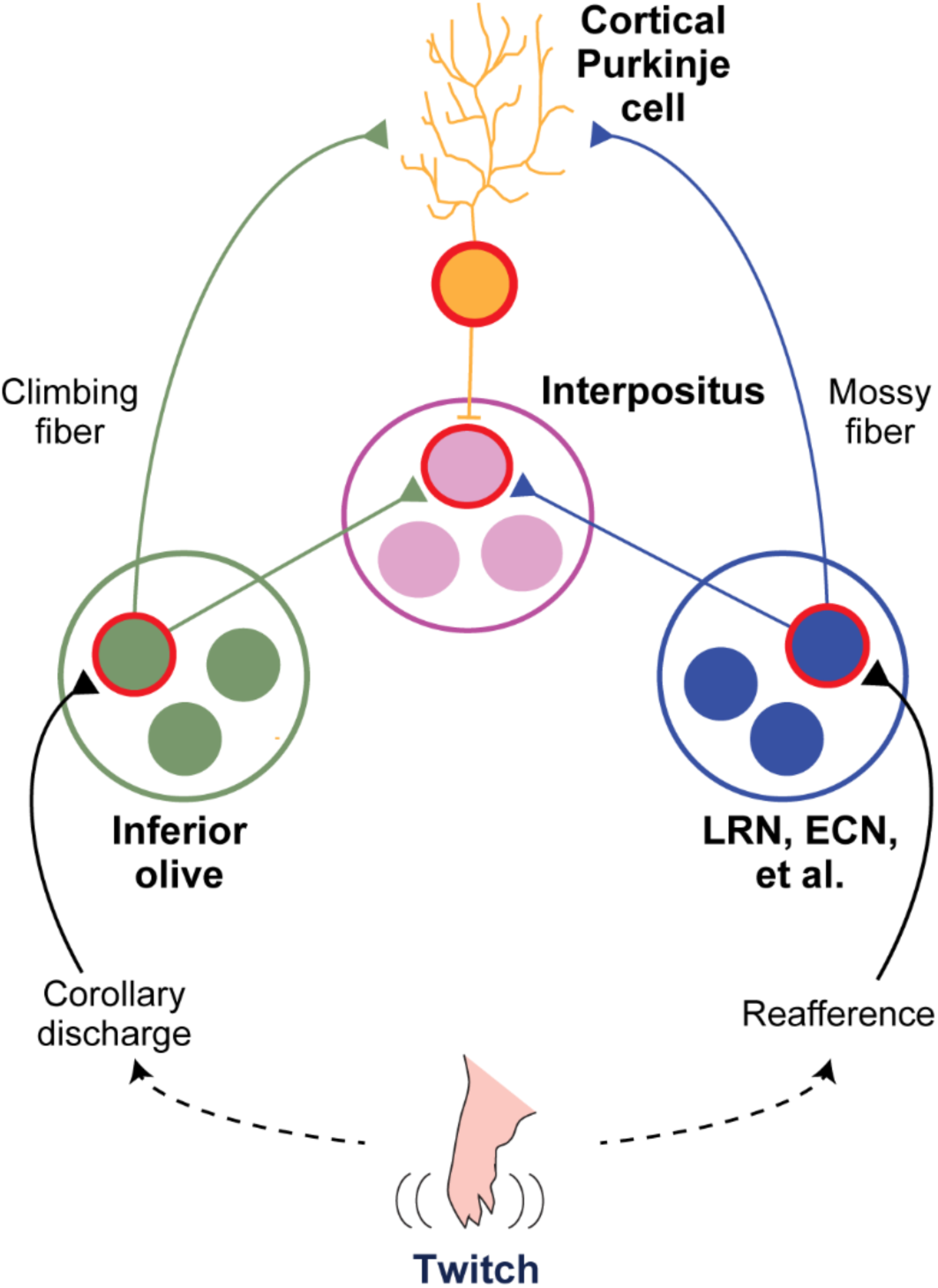
Developing a somatotopically organized closed-loop circuit in the cerebellar system. In early development, there are exuberant connections among key cerebellar and precerebellar components. Based on the current and previous findings, we propose that twitches, because they are associated with both corollary discharge and reafference, provide the opportunity for discrete activation of (for example) forelimb-associated neurons in the inferior olive and other precerebellar nuclei (e.g., LRN, lateral reticular nucleus; ECN, external cuneate nucleus). The resulting parallel coactivation of neurons in interpositus and cerebellar cortex enables the construction of somatotopically organized closed-loop circuits that will, over the next week, provide a foundation for the computation of internal models that allow for precise predictive control of movement.

### BCE: Before cerebellar error

The present results do not align with what is known about the adult cerebellar system. This lack of alignment is particularly clear with respect to the IO: In adults, it is understood that the IO provides teaching signals used in the computation of internal models. These teaching signals—also called “error signals”—convey input to the cerebellum that represents violations of expectation (Medina et al., 2002; Devor, 2002; Ito, 1984). Thus, whereas the adult IO readily responds to unexpected exafference (Wu et al., 2010; Rushmer et al., 1976; Gellman et al., 1985), it appears that the infant IO is driven primarily by corollary discharge (Mukherjee et al., 2018). Currently, we do not know when the IO becomes responsive to exafference, but we predict that it occurs before the onset of adult-like cerebellar function at P20 (Dooley et al, 2021). It also remains to be determined whether the IO’s responsivity to corollary discharge is exclusive to infancy or whether it persists into adulthood, perhaps still associated with twitching during sleep.

The functional relationship between the deep cerebellar nuclei and the IO also changes with age. In adults, IP projects back to IO to inhibit its activity and suppress expected stimuli (Hesslow & Ivarsson, 1996). Such inhibition is weak at P12 because the inhibitory connection from IP to IO is underdeveloped (Nicholson and Freeman, 2003). Thus, at an age when the cerebellar system lacks the capacity to compute predictive internal models (Dooley et al., 2021), the IO is not yet ready to detect error. These substantial age-related differences highlight the need to understand the infant’s cerebellar system within its appropriate developmental context. Just as primary motor cortex functions very differently in infants and adults (Dooley et al., 2018), we should not be surprised that the infant cerebellar system does not function identically to that of adults.

### Conclusions and Future Directions

Previous research from our lab documented twitch- and stimulation-related activity in precerebellar nuclei before P12 (Mukherjee et al., 2018; Tiriac and Blumberg, 2016), and the expression of a cerebellar-dependent internal model of movement in motor thalamus by P20 (Dooley et al., 2021). However, in our studies of neural activity within the cerebellum itself (Sokoloff et al., 2015b; Del Rio-Bermudez et al., 2016), we documented twitch-related activity but did not examine cerebellar responses to stimulation.

Thus, there was a gap in our understanding of how twitches—with their associated corollary discharge and reafference—might uniquely influence cerebellar development. The current study fills that gap by showing that at P12, neurons in IP exhibit robust responses to corollary discharge and reafference, but not exafference, indicating a prioritization of twitch-related input. This prioritization, coupled with the in-parallel activation of IP and cerebellar cortex (thus providing the opportunity for coincident activity in functionally related neurons), suggests a functional role for twitching in the development of somatotopically organized closed-loop circuits (Najac and Raman, 2017). Such circuits are necessary for the capacity to compute precise internal models.

Understanding the prioritization of reafference over exafference in the infant IP opens a host of new questions. For example, through what mechanism does twitch-related corollary discharge influence local processing within IP to enable the expression of reafference? Why is corollary discharge conveyed to IP via both the infant IO and LRN (Mukherjee et al., 2018)? In addition, understanding the functional development of the IO is critical to understanding how cerebellar internal models develop. For example, what is the mechanistic basis for why the adult IO, unlike the infant IO, readily responds to exafference? One possibility is that the development of inhibitory feedback from IP, beginning around P17, alters the IO’s responsivity to exafference (Nicholson and Freeman, 2003). Finally, questions remain concerning the necessity and sufficiency of twitches—as opposed to wake movements—for providing the corollary discharges that are needed to build internal models. Answering these and other questions will be necessary to fully explain the relative roles of self- and other-generated movements in the activity-dependent development of the cerebellar system.

We end by highlighting the predominance of sleep, particularly active sleep, in early mammalian and avian development (Jouvet-Mounier et al., 1970; Scriba et al., 2013; Roffwarg et al., 1966) and its contributions to the development of sensory and sensorimotor systems (Blumberg et al., 2022). One of active sleep’s most prominent features is twitches, whose unique characteristics make them particularly suitable for developing internal models (Dooley et al., 2021). Although twitches occur predominantly during active sleep in mammals and birds, spontaneous movements occur in other developing animals—for example, zebrafish (Wan et al., 2019) and flies (Carreira-Rosario et al., 2021; Zeng et al., 2021; Dilley et al., 2018; Shaw et al., 2000)—that, regardless of whether they exhibit active sleep per se, nonetheless sleep the most in early life (Kayser et al., 2014; Sorribes et al., 2013). Given that internal models are also exhibited by these animals (Markov et al., 2021; von Holst & Mittelstaedt, 1950) and the ubiquity of corollary discharge more generally across the animal kingdom (Crapse & Sommer, 2008), self-generated movements may provide a common mechanism for developing the capacity for self-guided, adaptive, and flexible movement.

## Acknowledgments

This research was supported by a grant from the National Institutes of Health (R37-HD081168) to M.S.B

